# Bayesian large-scale multiple regression with summary statistics from genome-wide association studies

**DOI:** 10.1101/042457

**Authors:** Xiang Zhu, Matthew Stephens

## Abstract

Bayesian methods for large-scale multiple regression provide attractive approaches to the analysis of genome-wide association studies (GWAS). For example, they can estimate heritability of complex traits, allowing for both polygenic and sparse models; and by incorporating external genomic data into the priors they can increase power and yield new biological insights. However, these methods require access to individual genotypes and phenotypes, which are often not easily available. Here we provide a framework for performing these analyses without individual-level data. Specifically, we introduce a “Regression with Summary Statistics” (RSS) likelihood, which relates the multiple regression coefficients to univariate regression results that are often easily available. The RSS likelihood requires estimates of correlations among covariates (SNPs), which also can be obtained from public databases. We perform Bayesian multiple regression analysis by combining the RSS likelihood with previously-proposed prior distributions, sampling posteriors by Markov chain Monte Carlo. In a wide range of simulations RSS performs similarly to analyses using the individual data, both for estimating heritability and detecting associations. We apply RSS to a GWAS of human height that contains 253,288 individuals typed at 1.06 million SNPs, for which analyses of individual-level data are practically impossible. Estimates of heritability (52%) are consistent with, but more precise, than previous results using subsets of these data. We also identify many previously-unreported loci that show evidence for association with height in our analyses. Software is available at https://github.com/stephenslab/rss.

## 1. Introduction

Consider the multiple linear regression model:

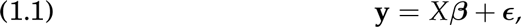

where **y** is an *n* × 1 (centered) vector, *X* is an *n × p* (column-centered) matrix, *β* is the *p* × 1 vector of multiple regression coefficients, and *E* is the error term. Assuming the “individual-level” data {*X*, **y**} are available, many methods exist to infer *β*. Here, motivated by applications in genetics, we assume that individual-level data are not available, but instead the summary statistics 
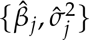
 from *p* simple linear regression are provided:

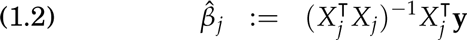

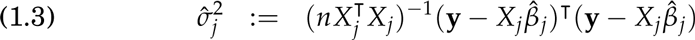

where *X_j_* is the *j* th column of *X*, *j* ∈ [*p*] := {1, …, *p*}. We also assume that information on the correlation structure among {*X_j_*} is available. With this in hand, we address the question: *how do we infer β using* 
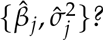
 Specifically, we derive a likelihood for *β* given 
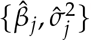
, and combine it with suitable priors to perform Bayesian inference for *β*.

This work is motivated by applications in genome-wide association studies (GWAS), which over the last decade have helped elucidate the genetics of dozens of complex traits and diseases [e.g. Donnelly (2008); McCarthy et al. (2008)]. GWAS come in various flavors – and can involve, for example, case-control data and/or related individuals – but here we focus on the simplest case of a quantitative trait (e.g. height) measured on random samples from a population. Model (1.1) applies naturally to this setting: the covariates *X* are the (centered) genotypes of *n* individuals at *p* genetic variants (typically Single Nucleotide Polymorphisms, or SNPs) in a study cohort; the response **y** is the quantitative trait whose relationship with genotype is being studied; and the coefficients *β* are the effects of each SNP on phenotype, estimation of which is a key inferential goal.

In GWAS individual-level data can be difficult to obtain. Indeed, for many publications no author had access to all the individual-level data. This is because many GWAS analyses involve multiple research groups pooling results across many cohorts to maximize sample size, and sharing individual-level data across groups is made difficult by many factors, including consent and privacy issues, and the substantial technical burden of data transfer, storage, management and harmonization. In contrast, summary data like 
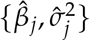
 are much easier to obtain: collaborating research groups often share such data to perform simple (though useful) “single-SNP” meta-analyses on a very large total sample size (Evangelou and Ioannidis, 2013). Furthermore these summary data are often made freely available on the Internet (Nature Genetics, 2012). In addition, information on the correlations among SNPs [referred to in population genetics as “linkage disequilibrium”, or LD; see Pritchard and Przeworski (2001)] is also available through public databases such as 1000 Genomes Project Consortium (2010). Thus, by providing methods for fitting the model (1.1) using only summary data and LD information, our work greatly facilitates the “multiple-SNP” analysis of GWAS data. For example, as we describe later, a single analyst (X.Z.) performed multiple-SNP analyses of GWAS data on adult height (Wood et al., 2014) involving 253,288 individuals typed at ~ 1.06 million SNPs, using modest computational resources (Section 6). Doing this for the individual-level data appears impractical.

Multiple-SNP analyses of GWAS compliment the standard single-SNP analyses in several ways. Multiple-SNP analyses are particularly helpful in fine-mapping causal loci, allowing for multiple causal variants in a region [e.g. Servin and Stephens (2007); Yang et al. (2012)]. In addition, they can increase power to identify associations [e.g. Hoggart et al. (2008); Guan and Stephens (2011)]; and can help estimate the overall proportion of phenotypic variation explained by genotyped SNPs (PVE; or “SNP heritability”) [e.g. Yang et al. (2010); Zhou, Carbonetto and Stephens (2013)]. See Sabatti (2013) and Guan and Wang (2013) for more extensive discussion. Despite these benefits, few GWAS are analyzed with multiple-SNP methods, presumably, at least in part, because existing methods require individual-level data that can be difficult to obtain. In addition, most multiple-SNP methods are computationally challenging for large studies [e.g. Peise, Fabregat-Traver and Bientinesi (2015); Loh et al. (2015)]. Our methods help with both these issues, allowing inference to be performed with summary-level data, and reducing computation by exploiting matrix bandedness (Wen and Stephens, 2010).

Because of the importance of this problem for GWAS, many recent publications have described analysis methods based on summary statistics. These include methods for estimation of effect size distribution (Park et al., 2010), conditional and joint multiple-SNP analysis (Yang et al., 2012), allelic heterogeneity detection (Ehret et al., 2012), single-SNP analysis with correlated phenotypes (Stephens, 2013) and heterogeneous subgroups (Wen and Stephens, 2014), gene-level testing of functional variants (Lee et al., 2015), joint analysis of functional genomic data and GWAS (Pickrell, 2014; Finucane et al., 2015), imputation of allele frequencies (Wen and Stephens, 2010) and single-SNP association statistics (Lee et al., 2013), fine mapping of causal variants (Hormozdiari et al., 2014; Chen et al., 2015), correction of inflated test statistics (Bulik-Sullivan et al., 2015), estimation of SNP heritability (Palla and Dudbridge, 2015), and prediction of polygenic risk scores (Vilhjalmsson et al., 2015). Together these methods adopt a variety of approaches, many of them tailored to their specific applications. Our approach, being based on a likelihood for the multiple regression coefficients *β*, provides the foundations for more generally-applicable methods. Having a likelihood opens the door to a wide range of statistical machinery for inference; here we illustrate this by using it to perform Bayesian inference for *β*, and specifically to estimate SNP heritability and detect associations.

Our work has close connections with recent Bayesian approaches to this problem, notably Hormozdiari et al. (2014) and Chen et al. (2015). These methods posit a model relating the observed *z*-scores {*β̂_j_/σ̂_j_*} to “non-centrality” parameters, and perform Bayesian inference on the noncentrality parameters. Here, we instead derive a likelihood for the regression coefficients *β* in (1.1), and perform Bayesian inference for *β*. These approaches are closely related, but working directly with *β* seems preferable to us. For example, the non-centrality parameters depend on sample size, which means that appropriate prior distributions may vary among studies depending on their sample size. In contrast, *β* maintains a consistent interpretation across studies. And working with *β* allows us to exploit previous work developing prior distributions for *β* for multiple-SNP analysis [e.g. Guan and Stephens (2011); Zhou, Carbonetto and Stephens (2013)]. We also give a more rigorous statement and derivation of the likelihood being used (Section 2.5), which provides insight into what approximations are being made and when they may be valid (Section 5). Finally, this previous work focused only on small genomic regions, whereas here we analyze whole chromosomes.

## 2. Likelihood based on summary data

We first introduce some notation. For any vector **v**, diag(**v**) denotes the diagonal matrix with diagonal elements **v**. Let 
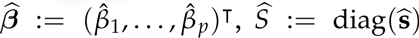
, and 
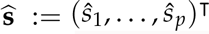
 where

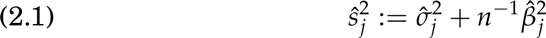

and 
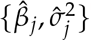
 are the single-SNP summary statistics (1.2, 1.3). We denote probability densities as *p* (·), and rely on the arguments to distinguish different distributions. Let 
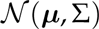
 denote the multivariate normal distribution with mean vector *μ* and covariance matrix Σ, and 
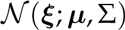
 denote its density at *ξ*.

In addition to the summary data 
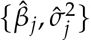
, we assume that we have an estimate, *R̂*, of the matrix *R* of LD (correlations) among SNPs in the population from which the genotypes were sampled. Typically *R̂* will come from some public database of genotypes in a suitable reference population; here, we use the shrinkage method from Wen and Stephens (2010) to obtain *R̂* from such a reference. The shrinkage method produces more accurate results than the sample correlation matrix (Section 4.1), and has the advantage that it produces a sparse, banded matrix *R̂*, which speeds computation for large genomic regions (Section 3.2). For our likelihood to be well-defined, *R̂* must be positive definite, and the shrinkage method also ensures this.

With this in place, the likelihood we propose for *β* is

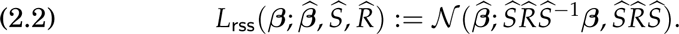

We refer to (2.2) as the “Regression with Summary Statistics” (RSS) likelihood. We provide a formal derivation in Section 2.5 (with proofs in Supplementary Appendix A), but informally the derivation assumes that i) the correlation of **y** with any single covariate (SNP) *X_j_* is small; and ii) the matrix *R* accurately reflects the correlation of the covariates (SNPs) in the population from which they were drawn.

The derivation of (2.2) also makes other assumptions that may not hold in practice: that all summary statistics are computed from the same samples, that there is no confounding due to population stratification (or that this has been adequately controlled for), and that genotypes used to computed summary statistics are accurate (so it ignores imputation error in imputed genotypes). Indeed, most analyses of individual-level data also make these last two assumptions. These assumptions can be relaxed, and generalizations of (2.2) derived; see Supplementary Appendix A. However these generalizations require additional information – beyond the basic single-SNP summary data (1.2, 1.3) – that is often not easily available. It is therefore tempting to apply (2.2) even when these assumptions may not hold. This is straightforward to do, but results in model misspecification and care is required; see Section 5.

### 2.1. Variations on RSS likelihood

We defined *Ŝ* by (2.1). In a GWAS context the sample sizes are often large and 
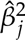
 are typically small (Table 1), and so 
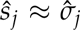
. Consequently, replacing 
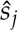
 in (2.2) with 
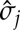
 produces a minor variation on the RSS likelihood that, for GWAS applications, differs negligibly from our definition (Supplementary Figure 4). This variation has slightly closer connections with existing work (Section 2.4).

**Table 1.** Summary of per-SNP sample squared correlation 
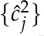
 and sample size {nj } for 42 large GWAS performed in European ancestry populations. The full names of phenotypes and references are provided in Supplementary Table 1. The summaries and histograms are across SNPs. The sample correlation ĉj between phenotype and SNP j is defined as 
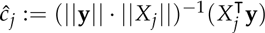
. Note that 
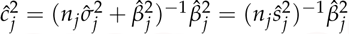
, and 
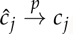
. The SD of sample sizes per SNP {nj } is NA when {nj } are not publicly available.

| GWAS phenotype | # of SNPs<br>(million) | $\log_{10}(\hat{c}_j^2)$ | | | | | Histogram | $\log_{10}(n)$ | | |
| --- | --- | --- | --- | --- | --- | --- | --- | --- | --- | --- |
|  |  | Min | Q1 | Median | Q3 | Max |  | Median | Mean | SD |
| Height (GIANT, 2010) | 2.82 | -12.64 | -6.25 | -5.60 | -5.12 | -2.90 |  | 5.26 | 5.26 | NA |
| Height (GIANT, 2014) | 2.53 | -10.74 | -6.06 | -5.41 | -4.93 | -2.54 |  | 5.40 | 5.37 | 0.09 |
| BMI (GIANT, 2015) | 2.54 | -10.89 | -6.30 | -5.65 | -5.18 | -2.66 |  | 5.37 | 5.34 | 0.09 |
| WHRadjBMI (GIANT, 2015) | 2.53 | -10.81 | -6.11 | -5.46 | -4.99 | -2.81 |  | 5.15 | 5.13 | 0.08 |
| HDL (GLGC, 2010) | 2.68 | -10.78 | -5.90 | -5.25 | -4.77 | -1.23 |  | 5.00 | 4.89 | 0.33 |
| HDL (GLGC, 2013) | 2.43 | -10.16 | -5.89 | -5.25 | -4.78 | -1.59 |  | 4.97 | 4.97 | 0.06 |
| LDL (GLGC, 2010) | 2.68 | -10.72 | -5.89 | -5.23 | -4.75 | -1.44 |  | 4.98 | 4.87 | 0.33 |
| LDL (GLGC, 2013) | 2.42 | -9.95 | -5.89 | -5.24 | -4.78 | -1.40 |  | 4.95 | 4.95 | 0.06 |
| TC (GLGC, 2010) | 2.68 | -10.33 | -5.91 | -5.25 | -4.77 | -1.38 |  | 5.00 | 4.89 | 0.33 |
| TC (GLGC, 2013) | 2.43 | -10.28 | -5.91 | -5.26 | -4.79 | -1.79 |  | 4.98 | 4.97 | 0.06 |
| TG (GLGC, 2010) | 2.68 | -10.55 | -5.89 | -5.24 | -4.76 | -1.17 |  | 4.98 | 4.87 | 0.33 |
| TG (GLGC, 2013) | 2.42 | -10.07 | -5.88 | -5.24 | -4.78 | -1.90 |  | 4.96 | 4.96 | 0.06 |
| Cigarettes per day (TAG, 2010) | 2.46 | -14.16 | -5.84 | -5.19 | -4.73 | -2.69 |  | 4.87 | 4.87 | NA |
| Smoking age of onset (TAG, 2010) | 2.43 | -11.27 | -5.82 | -5.18 | -4.73 | -3.44 |  | 4.87 | 4.87 | NA |
| Ever smoked (TAG, 2010) | 2.45 | -11.89 | -5.82 | -5.17 | -4.71 | -3.44 |  | 4.87 | 4.87 | NA |
| Former smoker (TAG, 2010) | 2.45 | -12.68 | -5.83 | -5.19 | -4.73 | -3.40 |  | 4.87 | 4.87 | NA |
| Years of education (SSGAC, 2013) | 2.14 | -7.10 | -5.70 | -5.30 | -4.85 | -3.51 |  | 5.10 | 5.10 | NA |
| College or not (SSGAC, 2013) | 2.25 | -8.37 | -5.93 | -5.33 | -4.88 | -3.39 |  | 5.10 | 5.10 | NA |
| Depressive (SSGAC, 2016) | 6.03 | -7.44 | -6.00 | -5.46 | -5.01 | -3.60 |  | 5.21 | 5.21 | NA |
| Neuroticism (SSGAC, 2016) | 6.04 | -7.88 | -5.92 | -5.45 | -4.98 | -3.29 |  | 5.23 | 5.23 | NA |
| Schizophrenia (PGC, 2014) | 9.43 | -12.50 | -6.01 | -5.35 | -4.88 | -3.04 |  | 5.18 | 5.18 | NA |
| Alzheimer (IGAP, 2013) | 7.04 | -11.20 | -5.69 | -5.04 | -4.57 | -1.33 |  | 4.73 | 4.73 | NA |
| CAD (CARDIoGRAM, 2011) | 2.42 | -17.43 | -5.84 | -5.18 | -4.71 | -2.74 |  | 4.91 | 4.88 | 0.08 |
| T2D (DIAGRAM, 2012) | 2.09 | -7.83 | -6.00 | -5.49 | -5.13 | -2.93 |  | 4.80 | 4.78 | 0.10 |
| Hb (HaemGen, 2012) | 2.58 | -9.79 | -5.64 | -4.99 | -4.52 | -2.47 |  | 4.74 | 4.68 | 0.15 |
| MCHC (HaemGen, 2012) | 2.57 | -9.72 | -5.62 | -4.98 | -4.51 | -2.50 |  | 4.70 | 4.65 | 0.15 |
| MCH (HaemGen, 2012) | 2.58 | -10.24 | -5.56 | -4.91 | -4.44 | -2.02 |  | 4.67 | 4.62 | 0.14 |
| MCV (HaemGen, 2012) | 2.59 | -11.02 | -5.61 | -4.96 | -4.48 | -2.09 |  | 4.71 | 4.66 | 0.15 |
| PCV (HaemGen, 2012) | 2.59 | -10.67 | -5.59 | -4.94 | -4.47 | -2.70 |  | 4.69 | 4.63 | 0.14 |
| RBC (HaemGen, 2012) | 2.56 | -8.92 | -5.55 | -4.91 | -4.45 | -2.11 |  | 4.69 | 4.63 | 0.15 |
| FGadjBMI (MAGIC, 2012) | 2.61 | -11.63 | -5.70 | -5.07 | -4.61 | -2.10 |  | 4.76 | 4.76 | NA |
| FIadjBMI (MAGIC, 2012) | 2.60 | -11.54 | -5.65 | -5.02 | -4.56 | -2.96 |  | 4.71 | 4.71 | NA |
| Heart rate (HRgene, 2013) | 2.52 | -12.08 | -5.88 | -5.23 | -4.76 | -2.88 |  | 4.95 | 4.93 | 0.07 |
| Serum urate (GUGC, 2013) | 2.44 | -10.36 | -5.95 | -5.30 | -4.83 | -1.49 |  | 5.04 | 5.03 | 0.02 |
| Gout (GUGC, 2013) | 2.54 | -12.17 | -5.80 | -5.15 | -4.69 | -2.68 |  | 4.84 | 4.84 | 0.01 |
| RA (Okada et al, 2014) | 7.70 | -8.16 | -5.44 | -4.97 | -4.55 | -1.09 |  | 4.77 | 4.77 | NA |
| IBD (IBDGC, 2015) | 12.70 | -13.05 | -5.49 | -4.84 | -4.38 | -2.07 |  | 4.54 | 4.54 | NA |
| CD (IBDGC, 2015) | 12.27 | -12.97 | -5.28 | -4.62 | -4.16 | -1.89 |  | 4.32 | 4.32 | NA |
| UC (IBDGC, 2015) | 12.24 | -12.69 | -5.40 | -4.75 | -4.28 | -2.07 |  | 4.44 | 4.44 | NA |
| CAD (CARDIoGRAM+C4D, 2015) | 9.46 | -15.00 | -6.26 | -5.61 | -5.14 | -2.62 |  | 5.27 | 5.27 | NA |
| MI (CARDIoGRAM+C4D, 2015) | 9.29 | -18.89 | -6.22 | -5.56 | -5.09 | -2.69 |  | 5.22 | 5.22 | NA |
| ANM (ReproGen, 2015) | 2.09 | -7.40 | -5.44 | -4.84 | -4.56 | -2.16 |  | 4.84 | 4.84 | NA |

Another variation comes from noting that the mean term in (2.2) does not change if we multiply *Ŝ* by any non-zero scalar constant: any constant will cancel out due to the presence of both *Ŝ* and 
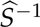
. Note further that 
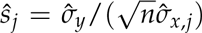
 where 
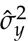
 is the sample variance of **y** (phenotype), and 
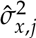
 the sample variance of *X_j_* (genotype at SNP *j*). Since *n* and 
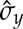
 are constants, the RSS likelihood is unchanged if we replaced *Ŝ* in the mean term with the matrix 
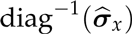
, where 
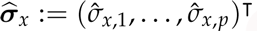
. That is:

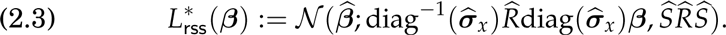

This variation on RSS helps emphasize the role of *Ŝ* in the mean term of (2.2): it is simply a convenience that exploits the fact that 
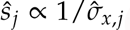
. The form (2.2) is more convenient in practice than (2.3), both because *Ŝ* is easily computed from commonly-used summary data and because the appearance of the same matrix *Ŝ* in the mean and variance terms of (2.2) produces algebraic simplifications that we exploit in our implementation. However, this convenient approach – which is also used in previous work (Section 2.4) – can contribute to model misspecification when, for example, different SNPs are typed on different samples; see Section 5.1.

### 2.2. Intuition

The RSS likelihood (2.2) is obtained by first deriving an approximation for *p*(*β̂* |*S*, *R*, *β*), where *S* is the diagonal matrix with the *j* th diagonal entry 
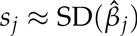
, of which *Ŝ* is an estimate (see Section 2.5 for details). Specifically, we have

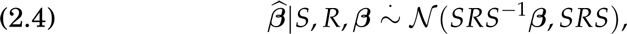

from which the RSS likelihood (2.2) is derived by plugging in the estimates {*Ŝ*, *R̂*} for {*S*, *R*}.

The distribution (2.4) captures three key features of the single-SNP association test statistics in GWAS. First, the mean of the single-SNP effect size estimate 
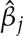
 depends on both its own effect and the effects of all SNPs that it “tags” (i.e. is highly correlated with):

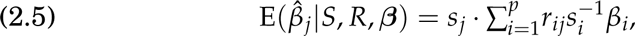

where *r_ij_* is the (*i*, *j*)-th entry of *R*. Second, the likelihood incorporates the fact that the estimated single-SNP effects are heteroscedastic:

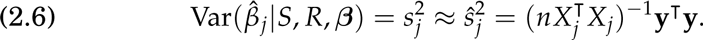

Since 
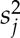
 is roughly proportional to 
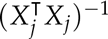
, the likelihood takes account of differences in the informativeness of SNPs due to their variation in allele frequency and imputation quality (Guan and Stephens, 2008). Third, single-SNP test statistics at SNPs in LD are correlated:

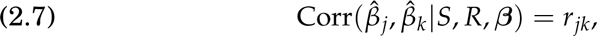

for any pair of SNP *j* and *k*.

Note that SNPs in LD with one another have “correlated” test statistics 
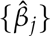
 for two distinct reasons. First, they share “signal”, which is captured in the mean term (2.5). This shared signal becomes a correlation if the true effects *β* are assumed to arise from some distribution and are then integrated out. Second, they share “noise”, which is captured in the correlation term (2.7). This latter correlation occurs even in the absence of signal (*β* = 0) and is due to the fact that the summary data are computed on the same samples. If the summary data were computed on independent sets of individuals, then this latter correlation would disappear (Section 5.1).

### 2.3. Connection with the full-data likelihood

When individual-level data are available the multiple regression model is

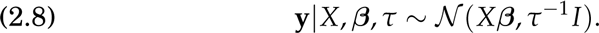

If we further assume the residual variance *τ*^−1^ is *known*, model (2.8) specifies a likelihood for *β*, which we denote *L*_mvn_(*β* ; **y**, *X*, *τ*). The following proposition gives conditions under which this full-data likelihood and RSS likelihood are equivalent.

#### Proposition 2.1.

Let 
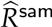
 denote the sample LD matrix computed from the genotypes *X* of the study cohort, 
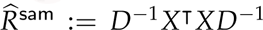
 where 
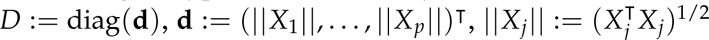

If *n > p*, *τ*^−1^ = *n* −1**y**T**y** and 
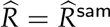
 then

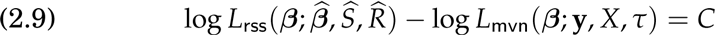

where *C* is some constant that does not depend on *β*.

The assumption *n > p* in Proposition 2.1 could possibly be relaxed, but certainly simplifies the proof. The key assumption then is *τ*^−1^ = *n* −1**y**T**y**; that is, that the total variance in **y** explained by *X* is negligible. This will typically not hold in a genome-wide context, but might hold, approximately, when fine mapping a small genomic region since SNPs in a small region typically explain a very small proportion of phenotypic variation^1^. Hence, provided that 
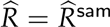
, RSS and its full-data counterpart will produce approximately the same inferential results in small regions. This is illustrated through simulations in Section 4.1 (Figure 1); see also Chen et al. (2015).

**Fig 1:**
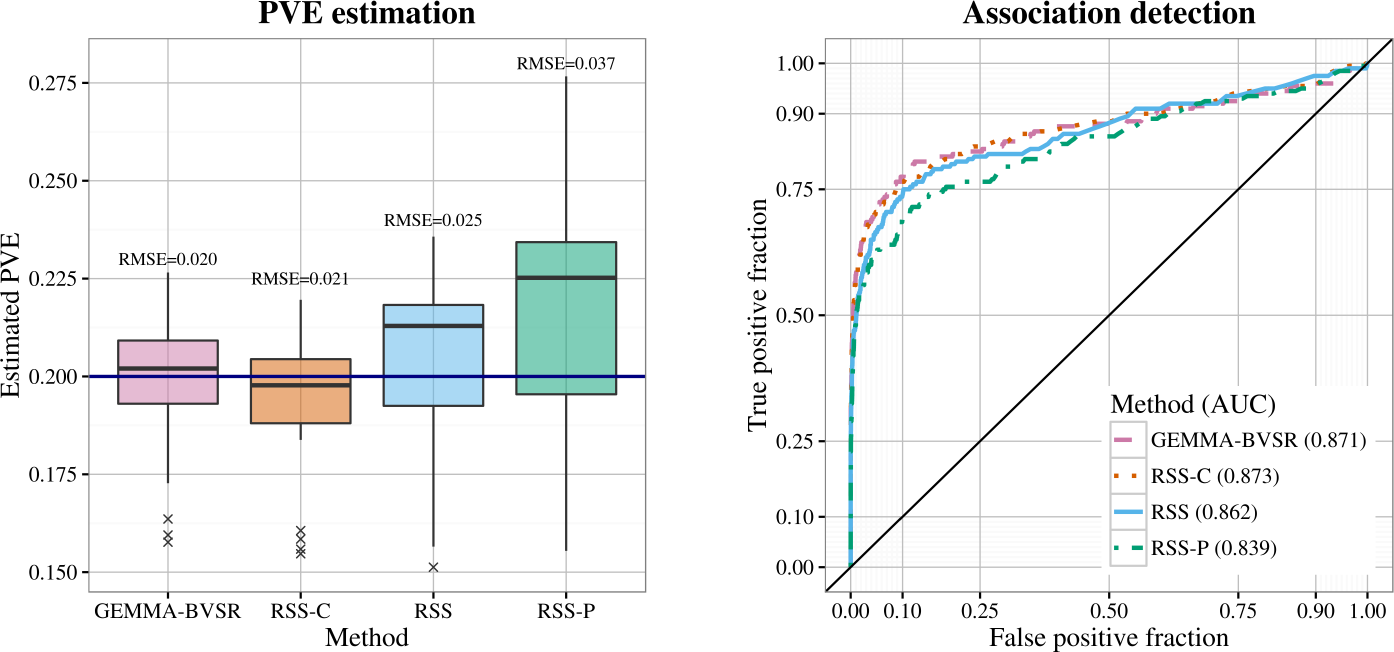
Comparison of PVE estimation and association detection on three types of LD matrix R: cohort sample LD (RSS-C), shrinkage panel sample LD (RSS) and panel sample LD (RSS-P). Performance of estimating PVE is measured by the root of mean square error (RMSE), where a lower value indicates better performance. Performance of detecting associations is measured by the area under the curve (AUC), where a higher value indicates better performance.

### 2.4. Connection with previous work

The RSS likelihood is connected to several previous approaches to inference from summary data, as we now describe. [These connections are precise for the variation on the RSS likelihood with 
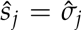
 (Section 2.1), which differs negligibly in practice from (2.2).]

In the simplest case, if *R̂* is an identity matrix, then 
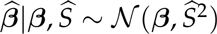
, which is the implied likelihood based on the standard confidence interval (Efron, 1993). Wakefield (2009) has recently popularized this likelihood for calculation of approximate Bayes factors; see also Stephens (2016).

If we let **z** denote the vector of single-SNP *z*-scores, 
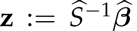
 and plug {*Ŝ*, *R̂*} into (2.4), then

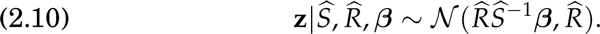

This is analogous to the likelihood proposed in Hormozdiari et al. (2014), 
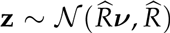
, where they refer to *ν* as the “non-centrality parameter”. If further *β* = 0, then 
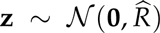
, a result that has been used for multiple testing adjustment [e.g. Seaman and Müller-Myhsok (2005); Lin (2005)], gene-based association detection [e.g. Liu et al. (2010)] and *z*-score imputation [e.g. Lee et al. (2013)].

If *β* is given a prior distribution that assumes zero mean and independence across all *j*, that is, 
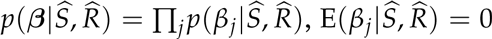
 then integrating *β* out in (2.10) yields 
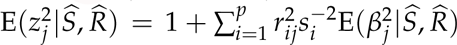
. This is a key element of LD score regression (Bulik-Sullivan et al., 2015); see Supplementary Appendix C for further details and discussion.

### 2.5. Derivation

We treat the (unobserved) genotypes of each individual, *x_i_* (the *i* th row of *X*), as being independent and identically distributed draws from some population. Without loss of generality, assume these have been centered, by subtracting the mean, so that E(*x_i_*) = 0. Let *σ_x,j_ >* 0 denote the population standard deviation (SD) of *x_ij_*, and *R* denote the *p* × *p* positive definite population correlation matrix, so Var(*x_i_*) := Σ*_x_* := diag(*σ_x_*) *R* diag(*σ_x_*), where *σ_x_* := (*σ*_*x*,1_, …, *σ_x,p_*)^T^.

We assume that the phenotypes **y** := (*y*_1_, …, *y_n_*)^T^ are generated from the multiple-SNP model (1.1), where E(*ɛ*) = 0 and Var(*ɛ*) = *τ*^−1^ *I_n_*. We also assume that *X*, *ɛ* and *β* are mutually independent.

Let **c** := (*c*_1_, …, *c_p_*)^T^ denote the vector of (population) correlations between the phenotype and each SNP:

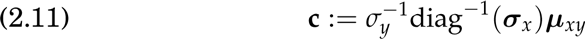

where *µ_xy_* := E(*x_i_y_i_*) and 
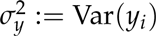
.

We first derive the asymptotic distribution of *β̂* (with *n* → ∞ and *p* fixed), using the Multivariate Central Limit Theorem and Delta Method.

#### Proposition 2.2.

Let 
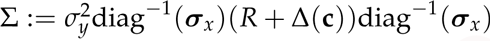
, where 
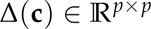
 is a continuous function of **c** and 
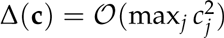
.

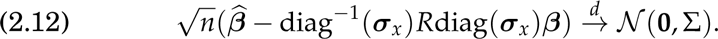

Proposition 2.2 suggests that the sampling distribution of *β̂* is close 
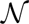
(diag^−1^(*σ_x_*)*R* diag(*σ_x_*)*β*, *n*^−1^Σ) for large *n*. Without additional assumptions, this may be the best^2^ probability statement that can be used to infer *β*. It is difficult to work with this asymptotic distribution, mainly because of the complicated form of Δ(**c**) (**Supplementary Appendix A**). However, we can justify ignoring this term in a typical GWAS by the fact that {*c_j_*} are typically small in GWAS (Table 1), and the following proposition:

#### Proposition 2.3.

Let 
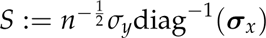
. For each 
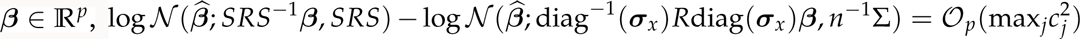
.

These propositions justify the approximate asymptotic distribution of *β̂* given in (2.4), provided *n* is large and 
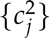
 close to zero, yielding

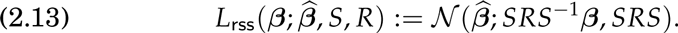

Finally, the RSS likelihood (2.2) is obtained by replacing the nuisance parameters {*S*, *R*} with their estimates {*Ŝ*, *R̂*}. There remains obvious potential for errors in the estimates *S*, *R* to impact inference, and we assess this impact empirically through simulations (Section **4**) and real data analyses (Section **6**).

## 3. Bayesian inference based on summary data

Using the RSS likelihood, we perform Bayesian inference for the multiple regression coefficients.

### 3.1. Prior specification

If {*S*, *R*} were known, then one could perform Bayesian inference by specifying a prior on *β* :

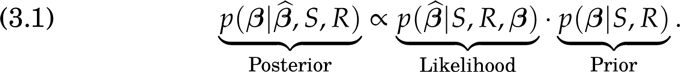

To deal with unknown {*S*, *R*} the RSS likelihood (**2.2**) approximates the likelihood in (**3.1**) by replacing {*S*, *R*} with their estimates {*S*, *R*}. We take a similar approach to prior specification: we specify a prior *p*(*β S*, *R*) and replace {*S*, *R*} with {*Ŝ*, *R̂*}.

Our prior specification is based on the prior from Zhou, Carbonetto and Stephens (2013) which was designed for analysis of individual-level GWAS data. This prior assumes that *β* is independent of *R a priori*, with the prior on *β_j_* being a mixture of two normal distributions

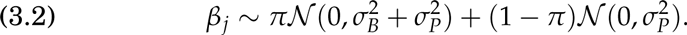

The motivation is that the first (“sparse”) component can capture rare “large” effects, while the second (“polygenic”) component can capture large numbers of very small effects. To specify priors on the variances 
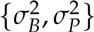
 Zhou, Carbonetto and Stephens (2013) introduce two free parameters *h*, *ρ* ∈ [0, 1] where *h* represents, roughly, the proportion of variance in **y** explained by *X*, and *ρ* represents the proportion of genetic variance explained by the sparse component. They write 
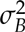
 and 
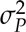
 as functions of *π*, *h*, *ρ* and place independent priors on the hyper-parameters (*π*, *h*, *ρ*):

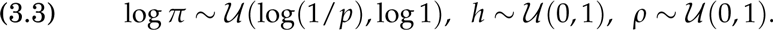

See Zhou, Carbonetto and Stephens (2013) for details.

Here we must modify this prior slightly because the original definitions of *σ_B_* and *σ_P_* depend on the genotypes *X* (which here are unknown) and the residual variance *τ*^−1^ (which does not appear in our likelihood). Specifically we define

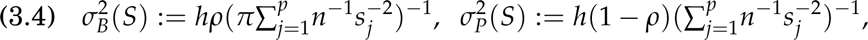

where *s_j_* is the *j* th diagonal entry of *S*. Because 
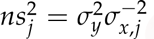
, definitions (3.4) ensure that the effect sizes of both components do not depend on *n*, and have the same measurement unit as the phenotype **y**. Further, with these definitions, *ρ* a nd *h* h ave i nterpretations s imilar t o t hose in previous work. Specifically, 
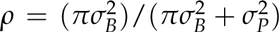
, so it represents the expected proportion of total genetic variation explained by the sparse components. Parameter *h* represents, roughly, the proportion of the total variation in **y** explained by *X*, as formalized by the following proposition:

#### Proposition 3.1.

If *β* |*S* is distributed as (3.2), with (3.4), then (3.5)

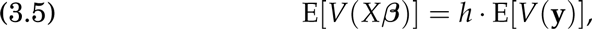

where *V*(*Xβ*) and *V*(**y**) are the sample variance of *Xβ* and **y** respectively.

Because of its similarity with the prior from “Bayesian sparse linear mixed model” [BSLMM, Zhou, Carbonetto and Stephens (2013)], we refer to our modified p rior a s B SLMM. We a lso i mplement a v ersion o f this prior where *ρ* = 1. This sets the polygenic variance 
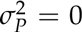
, making the prior on *β* sparse, and corresponds closely to the prior from “Bayesian variable selection regression” [BVSR, Guan and Stephens (2011)]. We therefore refer to this special case as BVSR here.

### 3.2. Posterior inference and computation

We use Markov chain Monte Carlo (MCMC) to sample from the posterior distribution of *β* ; see Supplementary Appendix B for details.

To fit t he R SS-BSLMM m odel, w e i mplement a n ew a lgorithm that is different from previous work (Zhou, Carbonetto and Stephens, 2013). Instead of integrating out *β* analytically, we perform MCMC sampling on *β* directly. Most of MCMC updates in this algorithm have linear complexity, with only a few “expensive” exceptions. The costs of these “expensive” updates are further reduced from being cubic in the total number of SNPs to being quadratic in this quantity, by leveraging the banded structure of LD matrix *R* (Wen and Stephens, 2010).

Our algorithm of fitting the RSS-BVSR model largely follows those developed in Guan and Stephens (2011), which exploit sparsity. Specifically, computation time per iteration scales cubically with the number of SNPs with non-zero effects, which is much smaller than the total number of SNPs under sparse assumptions. Setting a fixed maximum for the number of non-zero effects, and/or, using the banded LD structure to guide variable selection, can further improve computational performance, but we do not use these strategies here.

All computations were performed on a Linux system with a single Intel E5-2670 2.6GHz or AMD Opteron 6386 SE processor. Computation times for simulation studies and data analyses are shown in Supplementary Figure 5 and Supplementary Table 6 respectively. Software implementing the methods is available at https://github.com/stephenslab/rss.

Compared with existing summary-based methods, an important practical advantage of RSS is that multiple tasks can be performed using the same posterior sample of *β*. Here we focus on estimating PVE (SNP heritability) and detecting multiple-SNP associations.

#### 3.2.1. Estimating PVE

Given the full data {*X*, **y**} and the true value {*β*, *τ*} in model (1.1), Guan and Stephens (2011) define the PVE as

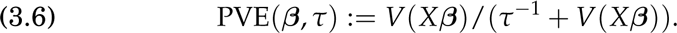

By this definition, PVE reflects the total proportion of sample phenotypic variation explained by available genotypes. Guan and Stephens (2011) then estimate PVE using the posterior sample of {*β*, *τ*}.

Because *X* is unknown here, we cannot compute PVE as defined above even if *β* and *τ* were known. Moreover, *τ* does not appear in our inference procedure. For these reasons we introduce the “Summary PVE” (SPVE) as an analogue of PVE for our setting:

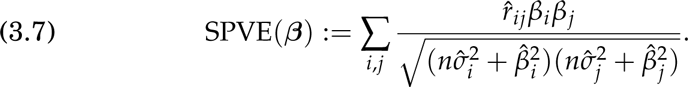

This definition is motivated by noting that PVE can be approximated by replacing *τ*^−1^ with V(**y**) − *V*(*Xβ*):

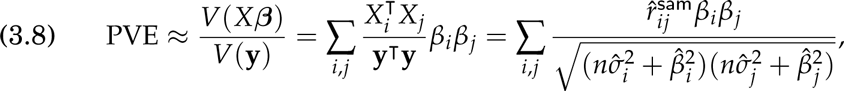

where 
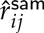
 is the (*i*, *j*)-entry of (unknown) sample LD matrix of the study cohort 
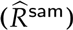
, which we approximate in SPVE by 
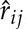
, and the last equation in (3.8) holds because of (1.2-1.3). Simulations using both synthetic and real genotypes show that SPVE is a highly accurate approximation to PVE, given the true value of *β* (Supplementary Figure 1).

We infer PVE using the posterior draws of SPVE, which are obtained by computing SPVE(*β*^(*i*)^) for each sampled value *β*^(*i*)^ from our MCMC algorithms. Unlike the original PVE (3.6), the definition of SPVE (3.7) is not bounded above by 1. Although we have not seen any estimates above 1 in our simulations or real data analyses, we expect this could occur if the posterior of *β* is poorly simulated and/or *R* is severely misspecified.

#### 3.2.2. Detecting genome-wide associations

Under the BVSR prior a natural summary of the evidence for a SNP being associated with phenotype is the posterior inclusion probability (PIP), Pr(*β_*j*_ ≠* 0 |**y**, *X*). Similarly, we define the PIP based on summary data

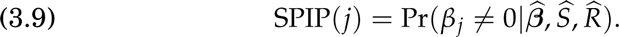

Here we estimate SPIP(*j*) by the proportion of MCMC draws for which *β_*j*_ ≠* 0. [We also provide a Rao-Blackwellised estimate in Supplementary Appendix B (Casella and Robert, 1996; Guan and Stephens, 2011).]

## 4. Simulations

We benchmark the RSS method through simulations, using real genotypes from Wellcome Trust Case Control Consortium (2007) (specifically, the *n* = 1458 individuals from the UK Blood Service Control Group) and simulated phenotypes. To reduce computation the simulations use genotypes from a single chromosome (12,758 SNPs on chromosome 16). One consequence of this is that the simulated effect sizes per SNP in some scenarios are often larger than would be expected in a typical GWAS (Table 1 and Supplementary Figure 3). This is, in some ways, not an ideal case for RSS, because the likelihood derivation assumes that effect sizes are small (Proposition 2.3). We use the simulations to i) investigate the effect of different choices for *R̂* ; and ii) demonstrate that inferences from RSS agree well with both the simulation ground truth, and with results from methods based on the full data [BVSR and BSLMM implemented in the software package GEMMA (Zhou and Stephens, 2012)].

### 4.1. Choice of LD matrix

The LD matrix *R̂* plays a key role in the RSS likelihood, as well as in previous work using summary data [e.g. Yang et al. (2012); Hormozdiari et al. (2014); Bulik-Sullivan et al. (2015)]. One simple choice for *R̂*, commonly used in previous work, is the sample LD matrix computed from a suitable “reference panel” that is deemed similar to the study population. This is a viable choice if the number of SNPs *p* is smaller than the number of individuals *m* in the reference panel, as the sample LD matrix is then invertible. However, for largescale genomic applications *p* ⪢ *m*, and the sample LD matrix is not invertible. Our proposed solution is to use the shrinkage estimator from Wen and Stephens (2010), which shrinks the off-diagonal entries of the sample LD matrix towards zero, resulting in an invertible matrix.

The shrinkage-based estimate of *R* can result in improved inference even if *p < m*. To illustrate this, we performed a small simulation study, with 982 SNPs within the ±5 Mb region surrounding the gene *IL27*. We simulated 20 independent datasets, each with 10 causal SNPs and PVE=0.2. (We also performed the simulations with the true PVE being 0.02 and 0.002; see Supplementary Figure 2.) For each dataset, we ran RSS-BVSR with two strategies for computing *R* from a reference panel (here, the 1480 control individuals in the WTCCC 1958 British Birth Cohort): the sample LD matrix (RSS-P), and the shrinkage-based estimate (RSS). We compared results with analyses using the full data (BVSR), and also with our RSS approach using the *cohort* LD matrix (RSS-C), which by Proposition 2.1 should produce results similar to the full data analysis. The results (Figure 1) show that using the shrinkage-based estimate for *R* produces consistently more accurate inferences – both for estimating PVE and detecting associations – than using the reference sample LD matrix, and indeed provides similar accuracy to the full data analysis.

### 4.2. Estimating PVE from summary data

Here we use simulations to assess the performance of RSS for estimating PVE. Using the WTCCC genotypes from 12,758 SNPs on chromosome 16, we simulated phenotypes under two genetic architectures:

- Scenario 1.1 (sparse): randomly select 50 “causal” SNPs, with effects ∼ 
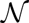
 (0, 1); effects of remaining SNPs are zero.
- Scenario 1.2 (polygenic): randomly select 50 “causal” SNPs, with effects ∼ 
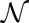
 (0, 1); effects of remaining SNPs are ∼ 
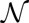
 (0, 0.001^2^).

For each scenario we simulated datasets with true PVE ranging from 0.05 to 0.5 (in steps of 0.05, with 50 independent replicates for each PVE). We ran RSS-BVSR on Scenario 1.1, and RSS-BSLMM on Scenario 1.2. Figure 2 summarizes the resulting PVE estimates. The estimated PVEs generally correspond well with the true values, but with a noticeable upward bias when the true PVE is large. We speculate that this upward bias is due to deviations from the assumption of small effects underlying RSS in Proposition 2.3. (Note that with 50 causal SNPs and PVE=0.5, on average each causal SNP explains 1% of the phenotypic variance, which is substantially higher than in typical GWAS; thus the upward bias in a typical GWAS may be less than in these simulations.)

**Fig 2:**
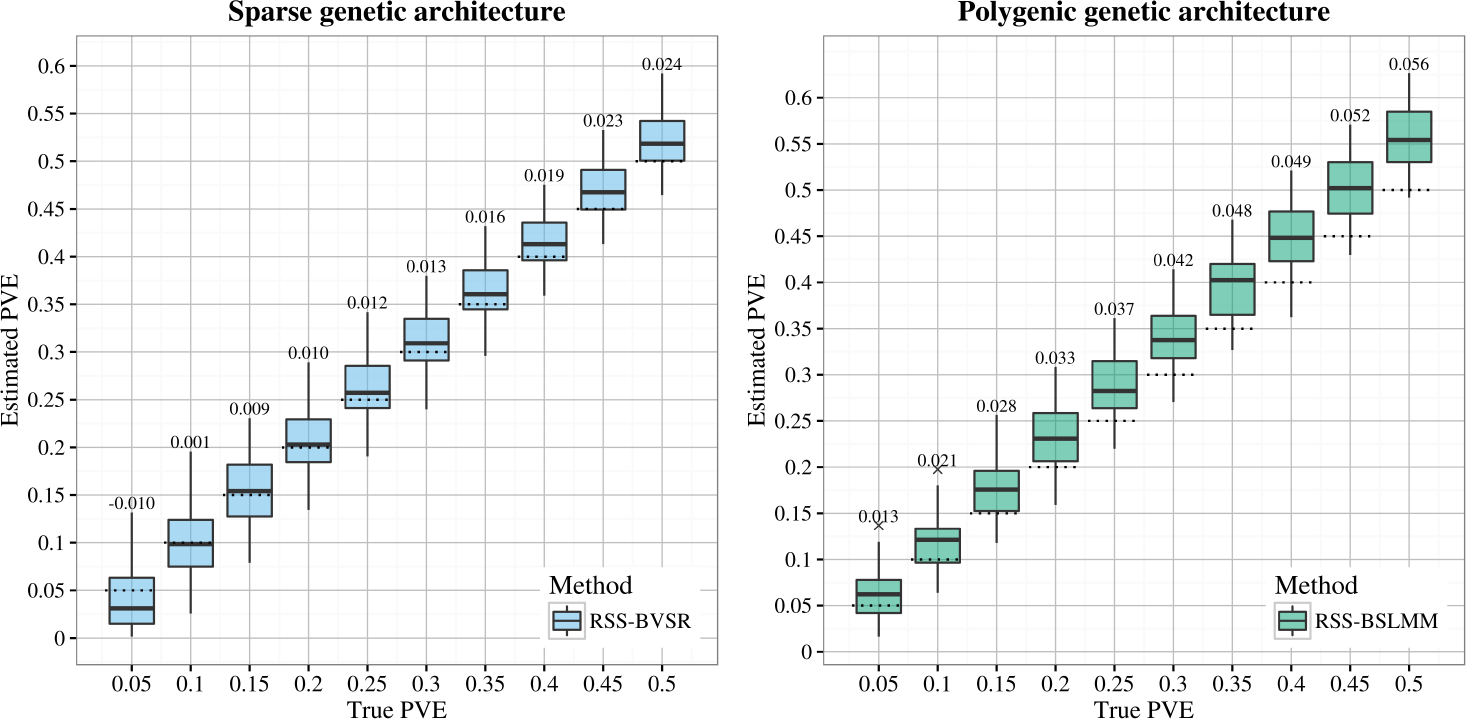
Comparison of true PVE with estimated PVE (posterior median) in Scenarios 1.1 (sparse) and 1.2 (polygenic). The dotted lines indicate the true PVEs, and the bias of estimates is reported on top of each box plot. Each box plot summarizes results from 50 replicates.

Next, we compare accuracy of PVE estimation using summary versus full data. With the genotype data as above we consider two scenarios:

- Scenario 2.1 (sparse): simulate a fixed number *T* of causal SNPs (*T* = 10, 100, 1000), with effect sizes coming from 
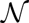
 (0, 1), and the effect sizes of the remaining SNPs are zero;
- Scenario 2.2 (polygenic): simulate two groups of causal SNPs, the first group containing a small number *T* of large-effect SNPs (*T* = 10, 100, 1000), plus another larger group of 10, 000 small-effect SNPs; the large effects are drawn from 
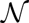
 (0, 1), the small effects are drawn from 
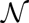
 (0, 0.001^2^), and the effect of the remaining SNPs are zero.

For each scenario we created datasets with true PVE 0.2 and 0.6 (20 independent replicates for each parameter combination). For Scenario 2.1 we compared results from the summary statistic methods (RSS-BVSR and RSS-BSLMM) with the corresponding full data methods (BVSR and BSLMM). For Scenario 2.2 we compared only the BSLMM methods, since the BVSR-based methods, which assume effects are sparse, are not well suited to this setting, in terms of both computation and accuracy (Zhou, Carbonetto and Stephens, 2013); see also Supplementary Appendix B. Figure 3 summarizes the results. With modest true PVE (0.2), BVSR and RSS-BVSR perform better than other methods when the true model is very sparse (e.g. Scenario 2.1, *T* = 10), whereas BSLMM and RSSBSLMM perform better when the true model is highly polygenic (e.g. Scenario 2.2, *T* = 1000). When the true PVE is large (0.6), the summary-based methods show an upward bias (Figure 3b and 3d), consistent with Figure 2. This bias is less severe when the true signals are more “diluted” (e.g. *T* = 1000), consistent with our speculation above that the bias is due to deviations from the “small effects” assumption. Overall, as expected, the summary data methods perform slightly less accurately than the full data methods. However, using different modeling assumptions (BVSR versus BSLMM) has a bigger impact on the results than using summary versus full data.

**Fig 3:**
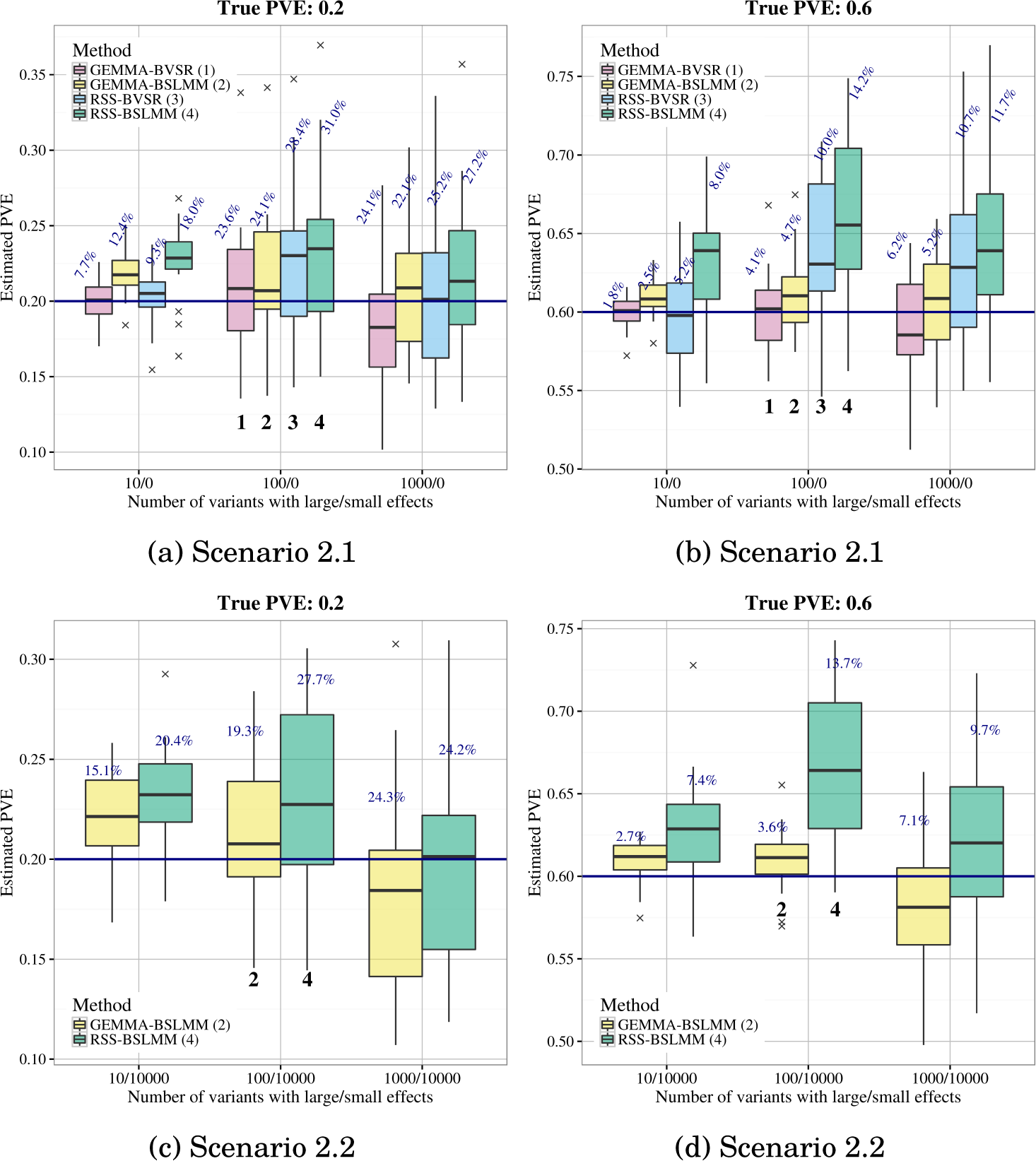
Comparison of PVE estimates (posterior median) from GEMMA and RSS in Scenario 2.1 and 2.2. The accuracy of estimation is measured by the relative RMSE, which is defined as the RMSE between the ratio of estimated over true PVEs and 1. Relative RMSE for each method is reported (percentages on top of box plots). The true PVEs are shown as the solid horizontal lines. Each box plot summarizes results from 20 replicates.

### 4.3. Power to detect associations from summary data

Previous studies using individual-level data have shown that multiple-SNP model can have higher power to detect SNP-phenotype associations than singleSNP analyses [e.g. Servin and Stephens (2007); Hoggart et al. (2008); Guan and Stephens (2011); Moser et al. (2015)]. Here we compare the power of multiple-SNP analyses based on summary data with those based on individual-level data. Specifically, we focus on comparing RSS-BVSR with BVSR, because the BVSR-based methods naturally select the associated SNPs (whereas BSLMM assumes that all SNPs are associated).

To compare associations detected by RSS-BVSR and BVSR, we simulated data under Scenario 2.1 above. With BVSR analyses, associations are most robustly assessed at the level of *regions* rather than at the level of individual SNPs (Guan and Stephens, 2011), so we compare the association signals from the two methods in sliding 200-kb windows (sliding each window 100kb at a time). Specifically, for each 200-kb region, and each method, we sum the PIPs of SNPs in the region to obtain the “Expected Number of included SNPs” (ENS), which summarizes the strength of association in that region. Results (Figure 4) show a strong correlation between the ENS values from the summary and individual data, across different numbers of causal variants and PVE values. Consequently, the summary data analyses have similar power to detect associations as the full data analyses (Figure 5). As above, the agreement of RSS-BVSR with BVSR is highest when PVE is diluted among many SNPs (e.g. *T* = 1000).

**Fig 4:**
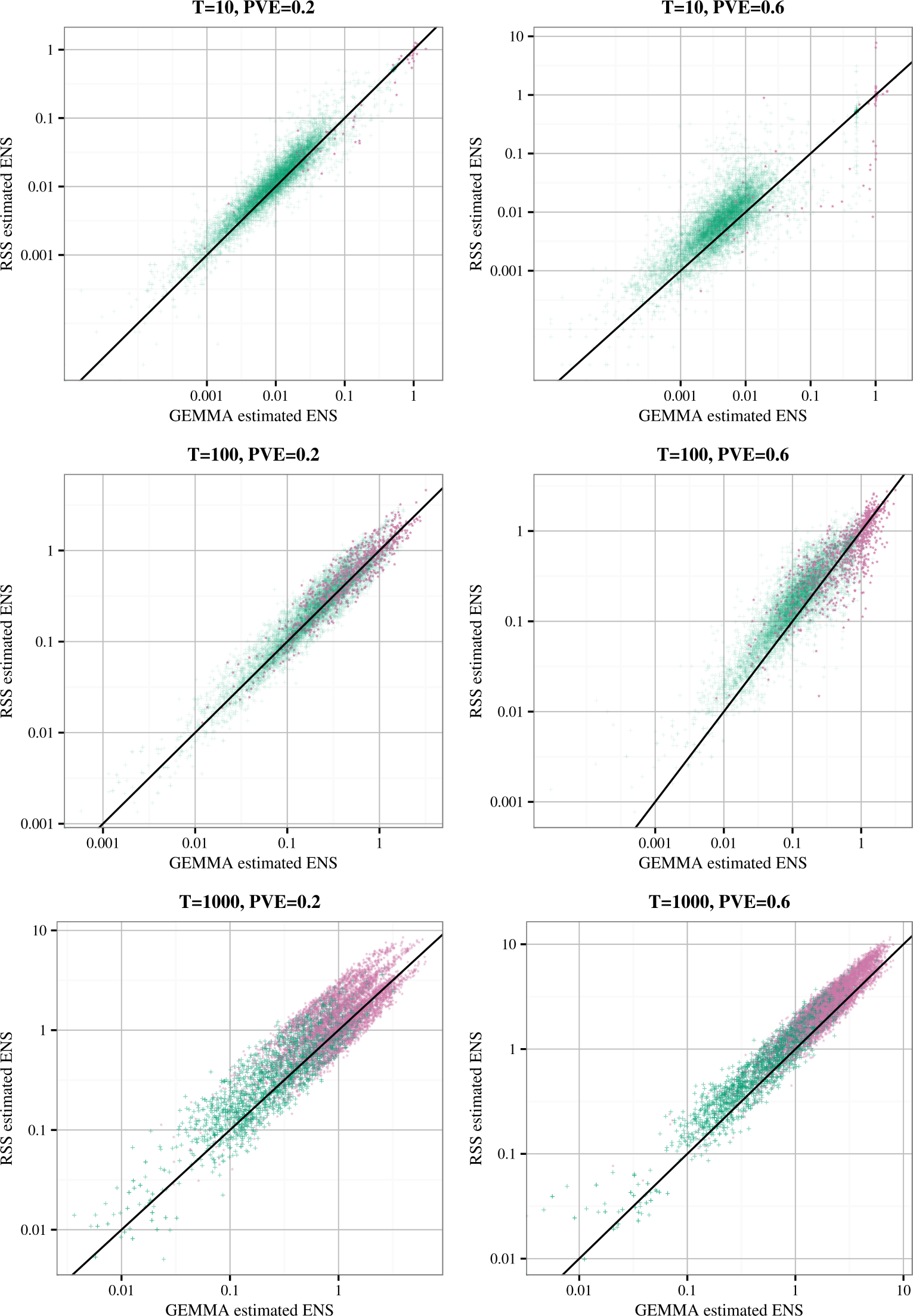
Comparison of the 200-kb region posterior expected numbers of included SNPs (ENS) for GEMMA-BVSR (x-axis) and RSS-BVSR (y-axis), based on the simulation study of Scenario 2.1. Each point is a 200-kb genomic region, colored according to whether it contains at least one causal SNP (reddish purple “*”) or not (bluish green “+”).

**Fig 5:**
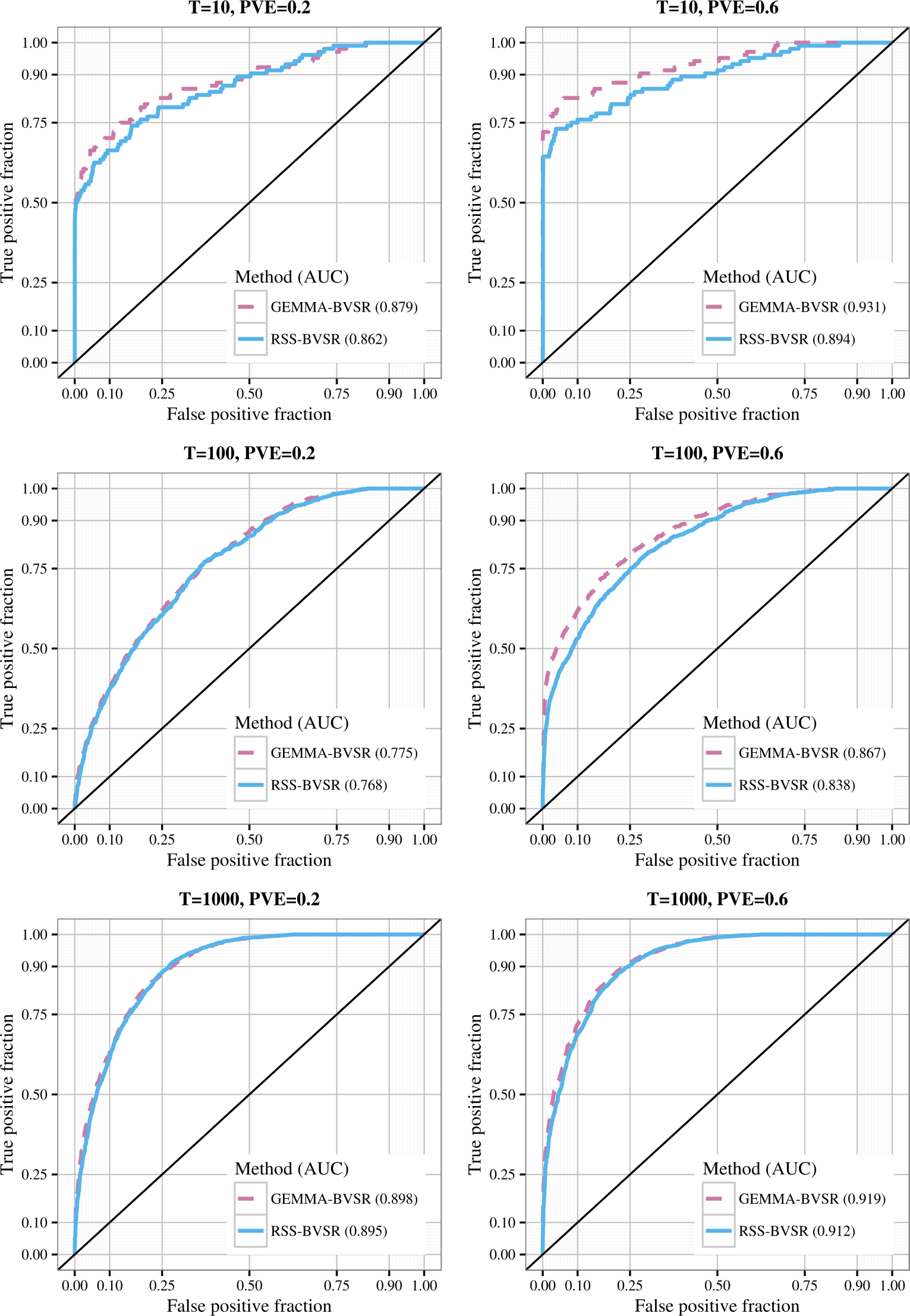
Trade-off between true and false positives for GEMMA-BVSR (reddish purple, dash) and RSS-BVSR (blue, solid) in simulations of Scenario 2.1.

## 5. Practical issues

The simulations in Section 4 show that, in idealized settings, RSS can largely recapitulate the results of a full multiple regression analysis. Specifically, these idealized settings involve summary data computed from a single set of individuals at fully observed genotypes. In practical applications summary data may deviate from this ideal. In addition, other issues, such as imputation quality and population stratification, can impact inferences from both full data and summary data. In this section we consider these practical issues, and make suggestions for how to deal with them both when generating summary data for distribution and when analyzing it.

### 5.1. Data on different individuals

In many studies data are available on different individuals at different SNPs (Table 1 and Supplementary Figure 7). This can happen for many reasons. For example, it can happen when combining information across individuals that are typed on different genotyping platforms. Or it can happen when combining data across multiple cohorts if quality control filters remove SNPs in some cohorts and not others.

It is important to note that the derivation of the RSS likelihood assumes that the summary statistics are generated from the same individuals at each SNP. Specifically, the covariances in likelihoods (2.2) and (2.3) depend on this assumption. [In contrast, the mean in likelihood (2.3) holds even if different individuals are used at each SNP; see Supplementary Appendix A for details.] To take an extreme example, if entirely different individuals are used to compute summary data for two SNPs then the correlation in their *β̂* values (given *β*) will be 0, even if the SNPs are in complete LD.

While RSS can be modified to allow for the use of different individuals when computing summary data at different SNPs [Propositions A.1 and A.2, Supplementary Appendix A; see also Zhang et al. (2016)], in practice this modification is unattractive because it requires considerable additional information in addition to the usual summary data – specifically, specification of sample overlaps for many pairs of SNPs. Instead, we recommend that genotype imputation [e.g. Servin and Stephens (2007); Marchini et al. (2007)] be used when generating GWAS summary data for public release, so that summary statistics are computed on the same individuals for each SNP.

When distributing summary data that are *not* computed on the same individuals, we recommend that at least the sample size used to compute data at each SNP also be made available, since these may be helpful both in modelling and in assessing the likely scope of the problem (Section 5.5). (Absent this, analysts may be able to estimate the number of individuals used at each SNP from *ŝ*_*j*_ and information on allele frequency of the SNP.)

### 5.2. Imputation quality

Many GWAS make use of genotype imputation to estimate genotypes that were not actually observed. Like almost all GWAS analysis methods that are used in practice, the RSS likelihood (2.2) does not formally incorporate the potential for error in the imputed genotypes.

In principle the RSS likelihood can be extended to account for imputation errors (Propositions A.3 and A.4, Supplementary Appendix A). However, this extension requires extra information – the imputation quality for each SNP – that is not always available. Fortunately, however, applying RSS to imputed genotypes, ignoring imputation quality, seems likely to provide sensible (if conservative) inferences in most cases. This is because imputation errors will tend to reduce estimated effects compared with what would have been obtained if all SNPs were typed: for example, if a SNP is poorly imputed then its estimated coefficient in the multiple regression model will be shrunk towards zero, and some of that SNP’s contribution to heritability will be lost. This issue is not restricted to RSS: indeed, it will also occur in analyses of individual-level data that use imputed genotypes.

A complimentary approach is to compile a list of SNPs that are expected, a priori, to be “well imputed” (Bulik-Sullivan et al., 2015), and to apply RSS only to these SNPs. This cannot remedy the loss of poorly-imputed SNPs’ contributions to heritability, but it may help avoid poorly-imputed SNPs undesirably influencing estimates of model hyper-parameters.

### 5.3. Population stratification

Another important issue that can impact many association studies is “confounding” due to population stratification (Devlin and Roeder, 1999; Price et al., 2010), which can cause over-estimation of genetic effects and heritability if not appropriately corrected for. A standard approach to dealing with this problem is to use methods such as principal components analysis (Price et al., 2006) and/or linear mixed models (Kang et al., 2010) to correct for stratification. These methods require access to the individual-level genotype data, and so cannot be used directly by analysts with access only to summary data. Instead they must be used by analysts who are computing the summary data for public distribution: doing so should substantially reduce the effects of confounding on summary data analyses, including RSS.

A complementary approach to dealing with population stratification is to directly model its effects on the summary data. One recent and innovative approach to this is LD score regression (Bulik-Sullivan et al., 2015), which uses the intercept of a regression of signal versus “LD score” to assess the effects of confounding. Along similar lines, we could modify the RSS likelihood to incorporate the effects of confounding by introducing an additional dispersion parameter (7.1); see Supplementary Appendix C. This modification would not require any additional summary data, and may have an additional benefit of improving robustness of RSS to other model misspecification issues (e.g. genotyping error, mismatches between LD in the reference panel and sample). However, this modification requires additional computation [some linear algebra simplifications we use when implementing (2.2) do not hold for (7.1)], and we have not yet implemented it.

### 5.4. Filtering and diagnostics

Some of the recommendations above can only be implemented when the summary data are being computed from individual data for public distribution, and not at a later stage when only the summary data are available. This raises the question, what can analysts with only access to summary data do to check that their results are likely reliable? This may be the trickiest part of summary data analysis: even with access to the full individual-level data it can be hard to assess all sources of bias and error. Recognizing that there is no universal approach that will guarantee reliable results we nonetheless hope to provide some useful suggestions.

Since the RSS likelihood (2.2) defines a statistical model, it is possible to perform a model fit diagnostic check. A generic approach to model checking (e.g. common in linear regression) is to first fit the model, compute residuals that measure deviations of observations from expected values, and then discard outlying observations before refitting the model. We have implemented an approach along these lines for identifying outlying SNPs, as follows. First, after fitting the model, we compute the residual (the difference between the observed *β̂* and its fitted expected value) at each SNP. We then perform a “leave-one-out” (LOO) check on each residual: we compute its conditional expectation and variance given the residuals at all other SNPs, and compute a diagnostic *z*-score based on how the observed residual compares with this expectation and variance. See Supplementary Appendix D for details. This approach targets SNPs whose summary data are most inconsistent with data at other near-by SNPs in LD. If the model is correctly specified for a given SNP then its diagnostic *z*-score approximately follows a standard normal distribution, from which a large deviation indicates potential misspecification. To assess robustness of RSS fit one can filter out SNPs with large diagnostic *z*-scores, and re-fit the RSS model on the remaining SNPs.

Other simpler filters are of course possible, and multiple filters can be used together. One widely-used filter simply discards SNPs with sample sizes lower than a certain cut-off (Pickrell, 2014). This can reduce problems caused by SNPs being typed on different subsets of individuals discussed above (Section 5.1). Another possibility is to filter out SNPs that are in very strong LD with one another, since these have potential for producing severe misspecification (Section 5.5). Some advantages of the model-based LOO diagnostic include that it could detect model misspecification problems from several sources – including genotyping error or misspecification of the LD matrix *R* – and not only those caused by typing of different individuals at different SNPs. Also, the sample size filter cannot be used unless the sample size for each SNP is made available, which is not always the case (Table 1). Finally, choice of threshold for the diagnostic *z*-score can be guided by the standard normal distribution; in contrast, selecting principled thresholds for sample sizes seems less straightforward (and a stringent threshold can yield conservative results; see Supplementary Figure 8). On the other hand, the LOO diagnostic may tend to filter out SNPs that show particularly strong signal (if they are not in LD with other SNPs), an undesirable property that should be remembered when interpreting results post-filtering (Supplementary Figure 9).

### 5.5. Extreme example

One way to help avoid problems with model misspecification is to be aware of the most severe ways in which things can go wrong. In this vein, we offer one illustrative example that we encountered when applying RSS to the summary data of a blood lipid GWAS (Global Lipids Genetics Consortium, 2013).

Table 2 shows summary statistics for high-density lipoprotein (HDL) cholesterol for seven SNPs in the gene *ADH5* that are in complete LD with one another in the reference panel (1000 Genomes European *r*^2^ = 1). If summary data were computed on the same set of individuals at each SNP, then they would be expected to vary very little among SNPs that are in such strong LD. And indeed, the RSS likelihood captures this expectation. However, in this case we see that the summary data actually vary considerably at some SNPs. The differences between one SNP (rs7683704) and the others are likely explained by the fact that this SNP was typed on more individuals: data at this SNP come from both GWAS (94,595 individuals) and Metabochip arrays (93,982 individuals). Thus this is an example of model misspecification due to SNPs being typed on different individuals. However, another SNP, rs13125919, also shows notable differences in summary data from the other SNPs, for reasons that are unclear to us. (This highlights a challenge of working with summary data – it is difficult to investigate the source of such anomalies without access to individual data.)

**Table 2.** Example of problems that can arise due to severe model misspecification. The table reports the sample sizes, single-SNP effect size estimates, SEs, and 1-SNP BFs of seven SNPs that are in complete LD in the reference panel (1000 Genomes, European ancestry). The 2-SNP BFs reported are for rs7683704 with each of the other SNPs. These unreasonably large 2-SNP BFs are due to model misspecification.

| SNP | $n_j$ | $\hat{\beta}_j$ | $\hat{\sigma}_j$ | 1-SNP $\log_{10}$ BF | 2-SNP $\log_{10}$ BF | $r^2$ |
| --- | --- | --- | --- | --- | --- | --- |
| rs7683704 | 187,124 | 0.0096 | 0.0058 | -0.676 | NA | 1.0 |
| rs13125919 | 94,311 | 0.0038 | 0.0079 | -1.084 | 172.638 | 1.0 |
| rs4699701 | 94,311 | 0.0054 | 0.0081 | -1.028 | 88.364 | 1.0 |
| rs17595424 | 94,274 | 0.0055 | 0.0081 | -1.024 | 83.925 | 1.0 |
| rs11547772 | 94,311 | 0.0056 | 0.0081 | -1.021 | 79.756 | 1.0 |
| rs7683802 | 94,311 | 0.0056 | 0.0081 | -1.021 | 79.756 | 1.0 |
| rs4699699 | 94,311 | 0.0058 | 0.0081 | -1.013 | 71.580 | 1.0 |

Whatever the reasons, applying RSS to these data results in severe model misspecification: based on their LD patterns RSS expects data at these SNPs to be almost identical, but they are not. This severe model misspecification can lead to unreliable results. For example, we used the RSS likelihood (2.2) to compute the 1-SNP and 2-SNP Bayes factors (BFs) [as in Servin and Stephens (2007); see also Chen et al. (2015)]. None of the SNPs shows evidence for marginal association with HDL (log10 1-SNP BF are all negative, indicating evidence for the null). However, the 2-SNP BFs for rs7683704 together with any of the other SNPs are unreasonably large, due to the severe model misspecification.

We emphasize that this is an extreme example, chosen to highlight the worst things that can go wrong. We did not come across any examples like this in spot-checks of results from the adult human height data below (Section 6). For simulations illustrating the effects of less extreme model mis-specification on PVE estimation see Supplementary Figure 6.

## 6. Analysis of summary data on adult height

We applied RSS to summary statistics from a GWAS of human adult height, involving 253,288 individuals of European ancestry typed at ~ 1.06 million SNPs (Wood et al., 2014). Accessing the individual-level genotypes and phenotypes would be a considerable undertaking; in contrast the summary data are easily and freely available^3^.

Following the protocol from Bulik-Sullivan et al. (2015), we filtered out poorly imputed SNPs and then removed SNPs absent from the genetic map of HapMap European-ancestry population Release 24 (Frazer et al., 2007). To avoid negative recombination rate estimates, we excluded SNPs in regions where the genome assembly had been rearranged. We also removed triallelic sites by manual inspection in BioMart (Smedley et al., 2015). This left 1,064,575 SNPs retained for analysis. We estimated the LD matrix *R* using phased haplotypes from 379 Europeanancestry individuals in 1000 Genomes Project Consortium (2010).

Although the summary data were generated after genotype imputation to the same reference panel [Section 1.1.2, Supplementary Note of Wood et al. (2014)], only 65% of the 1,064,575 analyzed SNPs were computed from the total sample (Supplementary Figure 7). This is because SNP filters applied by the consortium separately in each cohort often filtered out SNPs from a subset of cohorts [Section 1.1.4, Supplementary Note of Wood et al. (2014)]. As shown in Supplementary Appendix A, properly accounting for these non-overlapping samples would require sample overlap information that is not publicly available. Instead, we directly applied the original RSS likelihood (2.2) to the summary data. As discussed in Section 5.1, this simplification results in model misspecification. To assess the impact of this, in addition to the primary analysis using all the summary data, we also performed secondary analyses after applying the LOO residual diagnostic described in Section 5.4 to filter out SNPs whose diagnostic *z*-scores exceeded a threshold (2 or 3).

To reduce computation time and hardware requirement, we separately analyzed each of the 22 autosomal chromosomes so that all chromosomes were run in parallel in a computer cluster. In our analysis, each chromosome used a single CPU core. To assess convergence of the MCMC algorithm, we ran the algorithm on each dataset multiple times; results agreed well among runs (results not shown), suggesting no substantial problems with convergence. Here we report results from a single run on each chromosome with 2 million iterations. The CPU time of RSS-BVSR ranged from 1 to 36 hours, and the time of RSS-BSLMM ranged from 4 to 36 hours (Supplementary Table 6).

We first inferred PVE (SNP heritability) from these summary data. Figure 6 shows the estimated total and per-chromosome PVEs based on RSS-BVSR and RSS-BSLMM. For both methods, we can see an approximately linear relationship between PVE and chromosome length, consistent with a genetic architecture where many causal SNPs each contribute a small amount to PVE (a.k.a. “polygenicity”), and consistent with previous results using a mixed linear model (Yang et al., 2011) on three smaller individual-level datasets (number of SNPs: 593,521-687,398; sample size: 6,293-15,792). By summing PVE estimates across all 22 chromosomes, we estimated the total autosomal PVE to be 52.4%, with 95% credible interval [50.4%, 54.5%] using RSS-BVSR, and 52.1%, with 95% credible interval [50.3%, 53.9%] using RSS-BSLMM. Our estimates are consistent with, but more precise than, previous estimates based on individual-level data from subsets of this GWAS. Specifically, Wood et al. (2014) estimated PVE as 49.8%, with standard error 4.4%, from individual-level data of five cohorts (number of SNPs: 0.97-1.12 million; sample size: 1,145-5,668). The increased precision of the PVE estimates illustrates one benefit of being able to analyze summary data with a large sample size.

**Fig 6:**
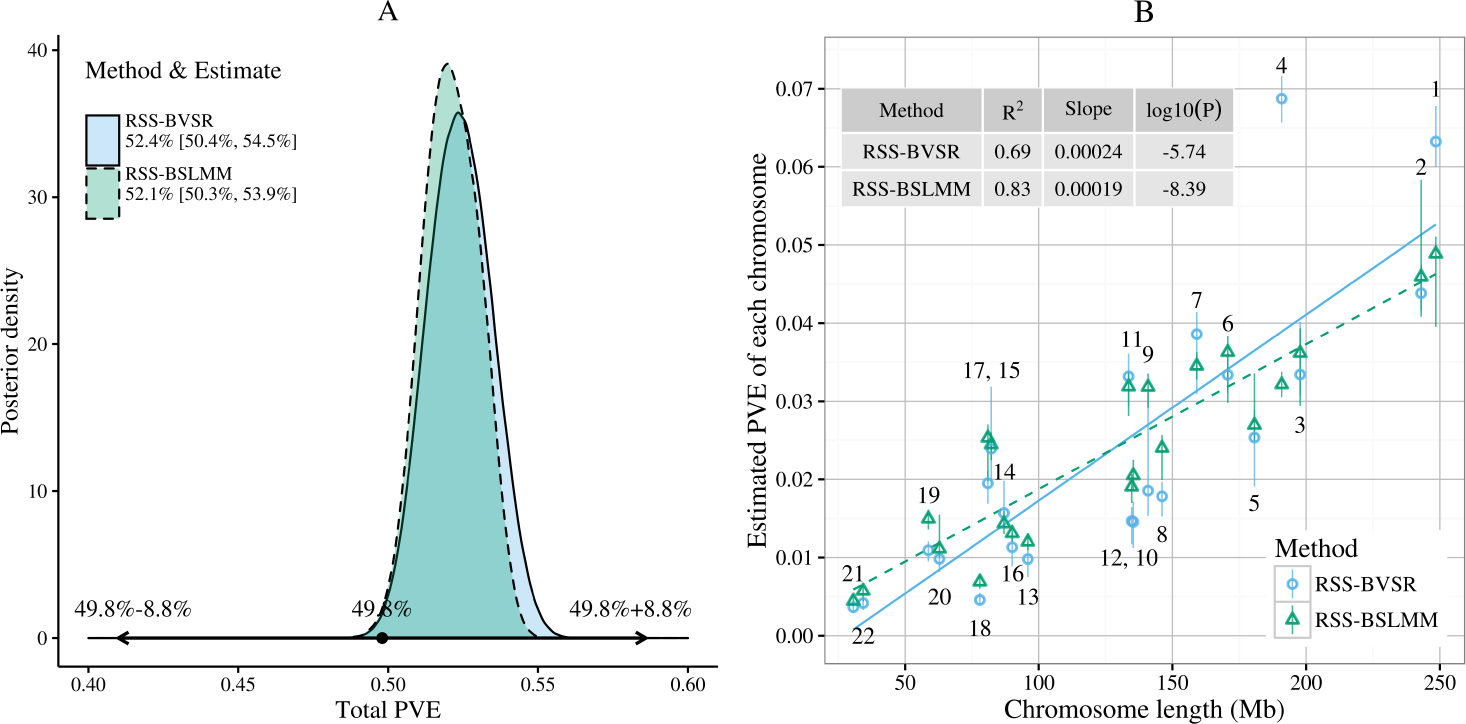
Posterior inference of PVE (SNP heritability) for adult human height. Panel A: posterior distributions of the total PVE, where the interval spanned by the arrows is the 95% confidence interval from Wood et al. (2014). Panel B: posterior median and 95% credible interval for PVE of each chromosome against the chromosome length, where each dot is labelled with chromosome number and the lines are fitted by simple linear regression (solid: RSS-BVSR; dash: RSSBSLMM). The regression output is shown in Supplementary Table 2. The data to reproduce Panel B are provided in Supplementary Table 3.

One caveat to these results is that the RSS likelihood (2.2) ignores confounding such as population stratification (Section 5.3). Here the summary data were generated using genomic control, principal components and linear mixed effects, to control for population stratification within each cohort [Section 1.1.3, Supplementary Note of Wood et al. (2014)]. Thus, we might hope that confounding has limited impact on PVE estimation. However, it is difficult to be sure that all confounding has been completely removed, and any remaining confounding could upwardly bias our estimated PVE. (Unremoved confounding could similarly bias estimates based on individual-level data.)

Next, we used RSS-BVSR to detect multiple-SNP associations, and compared results with previous analyses of these summary data. Using a stepwise selection strategy proposed by Yang et al. (2012), Wood et al. (2014) reported a total of 697 genome-wide significant SNPs (GWAS hits). Among them, 531 SNPs were within the ±40-kb regions with estimated ENS ≥ 1. Since only 384 GWAS hits were included in our filtered set of SNPs, we expected a higher replication rate for these included GWAS hits. Taking a region of ±40-kb around each of these 384 SNPs, our analysis identified almost all of these regions (371/384) as showing strong signal for association (estimated ENS ≥ 1). Only 125 of the 384 SNPs showed, individually, strong evidence for inclusion (estimated SPIP > 0.9). This suggests that, perhaps unsurprisingly, many of the reported associations are likely driven by a SNP in LD with the one identified in the original analysis.

To assess the potential for RSS to identify novel putative loci associated with human height, we estimated the ENS for ±40-kb windows across the whole genome. We identified 5194 regions with ENS ≥ 1, of which 2138 are putatively novel in that they are not near any of the previous 697 GWAS hits (distance > 1 Mb). Some of these 2138 regions are overlapping, but this nonetheless represents a large number of potential novel associations for further investigation. We manually examined the putatively novel regions with highest ENS, and identified several loci harboring genes that seem plausibly related to height. These include the gene *SCUBE1*, which is a co-receptor in promoting bone morphogenetic protein signalling (Liao, Tsao and Yang, 2016), the gene *WWOX*, which is a tumor suppressor linked to skeletal system morphogenesis (Del Mare et al., 2011; Aqeilan et al., 2008), the gene *IRX5*, which is essential for proximal and anterior skeletal formation (Li et al., 2014), and the gene *ALX1* (a.k.a. *CART1*), which is involved in bone development (Iioka et al., 2003). See Supplementary Table 5 for the full list of putatively new loci (estimated ENS > 3).

Finally, to check for misspecification we performed the LOO residual-based model diagnostic. Specifically, we ran the LOO residual imputation using the RSS-BVSR output, and then re-fitted the models on the filtered SNPs (absolute LOO *z*-score ≤ 2). This resulted in a substantial reduction in PVE estimates (RSS-BVSR: 34.0%, [32.9%, 35.0%]; RSSBSLMM: 45.3%, [44.7%, 46.0%]). However, this may reflect the fact that the filter removed 12% of SNPs, possibly biased towards SNPs showing associations (Supplementary Figure 9). By comparison, association results were more robust. Among the ±40-kb regions around the previous GWAS hits, our re-analyses identified 532 of the 697 total hits, and 373 of the 384 included hits. Moving the ±40-kb window across the genome, we identified 6426 regions with ENS e 1, of which 2798 were at least 1 Mb away from the 697 GWAS hits. Similar results based on filtering with a less stringent LOO threshold (3) are shown in Supplementary Table 4.

## 7. Discussion

We have presented a novel Bayesian method to infer multiple linear regression coefficients using simple linear regression summary statistics, and demonstrated its application in GWAS. On both simulated and real data our method produces results comparable to methods based on individual-level data. Compared with existing summary-based methods, our approach takes advantage of an explicit likelihood for the multiple regression coefficients, and thus provides a unified framework for various genome-wide analyses. We theoretically extend this framework to capture certain features of GWAS summary data, and provide practical suggestions when the theoretical extensions cannot be easily implemented. We illustrate the applications of our framework on heritability estimation and association detection. Other potential applications include training phenotype prediction models, prioritizing causal variants and testing gene effects.

We view the present work as the first stage of what could be done with RSS using GWAS summary statistics. One possibility for future work is to modify the RSS likelihood (2.2) to incorporate confounding by introducing an additional dispersion parameter *a* :

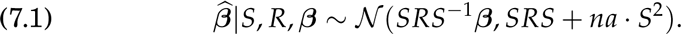

From model (7.1) we can derive relationships to LD score regression (Bulik-Sullivan et al., 2015), which distinguishes confounding biases from polygenicity using GWAS summary statistics; see Supplementary Appendix C for details. Another important extension is to integrate additional genomic information into the prior distributions. For instance, Carbonetto and Stephens (2013) allow the prior probability of each SNP being included to depend on a covariate, such as biological pathway membership,

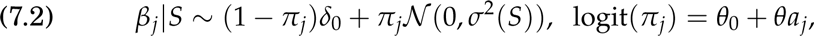

where *a_j_* = 1 if and only if SNP *j* is in the pathway. Unlike prior (3.2), prior (7.2) reflects that biologically related gene sets might preferentially harbor associated SNPs, essentially integrating the idea of gene set enrichment into GWAS (Wang, Li and Hakonarson, 2010). As a second example, some functional categories of the genome could contribute disproportionately to the heritability of complex traits (Gusev et al., 2014), which could be incorporated by letting the prior variance of the SNP effects depend on functional categorization, for example by

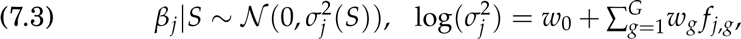

where *f_j,g_* = 1 when SNP *j* belongs to category *g*, *w*_0_ captures the baseline (log) heritability and {*w_g_*} reflect the contribution of each category. This could provide a different way to partition heritability by functional annotation using GWAS summary statistics (Finucane et al., 2015).

## Acknowledgements

We thank the Editor, Associate Editor and two anonymous referees for their constructive comments. We thank Xin He, Rina Foygel Barber, Peter Carbonetto, Yongtao Guan, Xiaoquan Wen and Xiang Zhou for helpful discussions. We thank Raman Shah and John Zekos for technical support. This study makes use of data generated by the Wellcome Trust Case Control Consortium. A full list of the investigators who contributed to the generation of the data is available from www.wtccc.org.uk. Data on adult human height have been contributed by investigators of the Genetic Investigation of Anthropometric Traits (GIANT) consortium. This work was completed in part with resources provided by the University of Chicago Research Computing Center.

## Footnotes

1 There are exceptions; for example the human leukocyte antigen (HLA) region is estimated to explain 11-37% of the heritability of rheumatoid arthritis (Kurkó et al., 2013)

2 A more rigorous approximation of likelihood based on the convergence in distribution requires additional technical assumptions; see **Boos (1985)** and **Sweeting (1986)**

3 https://www.broadinstitute.org/collaboration/giant/index.php/GIANT_consortium_data_files

## References

Aqeilan, R. I., Hassan, M. Q., de Bruin, A., Hagan, J. P., Volinia, S., Palumbo, T., Hus-sain, S., Lee, S.-H., Gaur, T., Stein, G. S. et al. (2008). The WWOX tumor suppressor is essential for postnatal survival and normal bone metabolism. Journal of Biological Chemistry 283 21629–21639.

Boos, D. D. (1985). A converse to Scheffe’s Theorem. The Annals of Statistics 13 423–427.

Bulik-Sullivan, B., Loh, P.-R., Finucane, H., Ripke, S., Yang, J., Psychiatric Genomics Consortium, Schizophrenia Working Group, Patterson, N., Daly, M. J., Price, A. L. and Neale, B. M. (2015). LD score regression distinguishes confounding from polygenicity in genome-wide association studies. Nature Genetics 47 291–295.

Carbonetto, P. and Stephens, M. (2013). Integrated enrichment analysis of variants and pathways in genome-wide association studies indicates central role for IL-2 Signaling genes in Type 1 Diabetes, and Cytokine Signaling genes in Crohn’s Disease. PLoS Genetics 9 e1003770.

Casella, G. and Robert, C. P. (1996). Rao-Blackwellisation of sampling schemes. Biometrika 83 81–94.

Chen, W., Larrabee, B. R., Ovsyannikova, I. G., Kennedy, R. B., Haralambieva, I. H., Poland, G. A. and Schaid, D. J. (2015). Fine mapping causal variants with an approximate Bayesian method using marginal test statistics. Genetics 200 719–736.

Wellcome Trust Case Control Consortium (2007). Genome-wide association study of 14,000 cases of seven common diseases and 3,000 shared controls. Nature 447 661–678.

1000 Genomes Project Consortium (2010). A map of human genome variation from population-scale sequencing. Nature 467 1061–1073.

Global Lipids Genetics Consortium (2013). Discovery and refinement of loci associated with lipids levels. Nature Genetics 45 1274–1283.

Del Mare, S., Kurek, K. C., Stein, G. S., Lian, J. B. and Aqeilan, R. I. (2011). Role of the WWOX tumor suppressor gene in bone homeostasis and the pathogenesis of osteosarcoma. American Journal of Cancer Research 1 585.

Devlin, B. and Roeder, K. (1999). Genomic control for association studies. Biometrics 55 997–1004.

Donnelly, P. (2008). Progress and challenges in genome-wide association studies in humans. Nature 456 728–731.

Efron, B. (1993). Bayes and likelihood calculations from confidence intervals. Biometrika 80 3–26.

Ehret, G. B., Lamparter, D., Hoggart, C. J., Whittaker, J. C., Beckmann, J. S., Kutalik, Z., of Anthropometric Traits Consortium, G. I. et al. (2012). A multi-SNP locus-association method reveals a substantial fraction of the missing heritability. The American Journal of Human Genetics 91 863–871.

Evangelou, E. and Ioannidis, J. P. (2013). Meta-analysis methods for genome-wide association studies and beyond. Nature Reviews Genetics 14 379–389.

Finucane, H. K., Bulik-Sullivan, B., Gusev, A., Trynka, G., Reshef, Y., Loh, P.-R., Anttila, V., Xu, H., Zang, C., Farh, K. et al. (2015). Partitioning heritability by functional annotation using genome-wide association summary statistics. Nature Genetics.

Frazer, K. A., Ballinger, D. G., Cox, D. R., Hinds, D. A., Stuve, L. L., Gibbs, R. A., Belmont, J. W., Boudreau, A., Hardenbol, P., Leal, S. M. et al. (2007). A second generation human haplotype map of over 3.1 million SNPs. Nature 449 851–861.

Nature Genetics (2012). Asking for more. Nature Genetics 44 733.

Guan, Y. and Stephens, M. (2008). Practical issues in imputation-based association mapping. PLoS Genetics 4 e1000279.

Guan, Y. and Stephens, M. (2011). Bayesian variable selection regression for genomewide association studies, and other large-scale problems. The Annals of Applied Statistics 5 1780–1815.

Guan, Y. and Wang, K. (2013). Whole-genome multi-SNP-phenotype association analysis. In Advances in Statistical Bioinformatics (K.-A. Do, Z. S. Qin and M. Vannucci, eds.) 224–243. Cambridge University Press.

Gusev, A., Lee, S. H., Trynka, G., Finucane, H., Vilhjálmsson, B. J., Xu, H., Zang, C., Ripke, S., Bulik-Sullivan, B., Stahl, E., Kähler, A. K., Hultman, C. M., Purcell, S. M., Mccarroll, S. A., Daly, M., Pasaniuc, B., Sullivan, P. F., Neale, B. M., Wray, N. R., Raychaudhuri, S. and Price, A. L. (2014). Partitioning heritability of regulatory and cell-type-specific variants across 11 common diseases. The American Journal of Human Genetics 95 535–552.

Hoggart, C. J., Whittaker, J. C., De Iorio, M. and Balding, D. J. (2008). Simultaneous analysis of all SNPs in genome-wide and re-sequencing association studies. PLoS Genetics 4 e1000130.

Hormozdiari, F., Kostem, E., Kang, E. Y., Pasaniuc, B. and Eskin, E. (2014). Identifying causal variants at loci with multiple signals of association. Genetics 198 497–508.

Iioka, T., Furukawa, K., Yamaguchi, A., Shindo, H., Yamashita, S. and Tsukazaki, T. (2003). P300/CBP acts as a coactivator to cartilage homeoprotein-1 (Cart1), paired like homeoprotein, through acetylation of the conserved lysine residue adjacent to the homeodomain. Journal of Bone and Mineral Research 18 1419–1429.

Kang, H. M., Sul, J. H., Service, S. K., Zaitlen, N. A., Kong, S.-y., Freimer, N. B., Sabatti, C., Eskin, E. et al. (2010). Variance component model to account for sample structure in genome-wide association studies. Nature Genetics 42 348–354.

Kurkó, J., Besenyei, T., Laki, J., Glant, T. T., Mikecz, K. and Szekanecz, Z. (2013). Genetics of rheumatoid arthritis–a comprehensive review. Clinical reviews in allergy & immunology 45 170–179.

Lee, D., Bigdeli, T. B., Riley, B. P., Fanous, A. H. and Bacanu, S.-A. (2013). DIST: direct imputation of summary statistics for unmeasured SNPs. Bioinformatics 29 2925–2927.

Lee, D., Williamson, V. S., Bigdeli, T. B., Riley, B. P., Fanous, A. H., Vladimirov, V. I. and Bacanu, S.-A. (2015). JEPEG: a summary statistics based tool for gene-level joint testing of functional variants. Bioinformatics 31 1176–1182.

Li, D., Sakuma, R., Vakili, N. A., Mo, R., Puviindran, V., Deimling, S., Zhang, X., Hopyan, S. and Hui, C.-c. (2014). Formation of proximal and anterior limb skeleton requires early function of Irx3 and Irx5 and is negatively regulated by Shh signaling. Developmental cell 29 233–240.

Liao, W.-J., Tsao, K.-C. and Yang, R.-B. (2016). Electrostatics and N-glycan-mediated membrane tethering of *SCUBE1* is critical for promoting bone morphogenetic protein signalling. Biochemical Journal 473 661–672.

Lin, D. (2005). An efficient Monte Carlo approach to assessing statistical significance in genomic studies. Bioinformatics 21 781–787.

Liu, J. Z., Mcrae, A. F., Nyholt, D. R., Medland, S. E., Wray, N. R., Brown, K. M., Hayward, N. K., Montgomery, G. W., Visscher, P. M., Martin, N. G. et al. (2010). A versatile gene-based test for genome-wide association studies. The American Journal of Human Genetics 87 139–145.

Loh, P.-R., Tucker, G., Bulik-Sullivan, B. K., Vilhjalmsson, B. J., Finucane, H. K., Chasman, D. I., Ridker, P. M., Neale, B. M., Berger, B., Patterson, N. et al. (2015). Efficient Bayesian mixed model analysis increases association power in large cohorts. Nature Genetics 47 284–290.

Marchini, J., Howie, B., Myers, S., McVean, G. and Donnelly, P. (2007). A new multipoint method for genome-wide association studies by imputation of genotypes. Nature Genetics 39 906–913.

Mccarthy, M. I., Abecasis, G. R., Cardon, L. R., Goldstein, D. B., Little, J., Ioannidis, J. P. and Hirschhorn, J. N. (2008). Genome-wide association studies for complex traits: consensus, uncertainty and challenges. Nature Reviews Genetics 9 356–369.

Moser, G., Lee, S. H., Hayes, B. J., Goddard, M. E., Wray, N. R. and Visscher, P. M. (2015). Simultaneous discovery, estimation and prediction analysis of complex traits using a Bayesian mixture model. PLoS Genetics 11 e1004969.

Palla, L. and Dudbridge, F. (2015). A fast method that uses polygenic scores to estimate the variance explained by genome-wide marker panels and the proportion of variants affecting a trait. The American Journal of Human Genetics 97 250–259.

Park, J.-H., Wacholder, S., Gail, M. H., Peters, U., Jacobs, K. B., Chanock, S. J. and Chatterjee, N. (2010). Estimation of effect size distribution from genome-wide association studies and implications for future discoveries. Nature Genetics 42 570–575.

Peise, E., Fabregat-Traver, D. and Bientinesi, P. (2015). High performance solutions for big-data GWAS. Parallel Computing 42 75–87.

Pickrell, J. K. (2014). Joint analysis of functional genomic data and genome-wide association studies of 18 human traits. The American Journal of Human Genetics 94 559–573.

Price, A. L., Patterson, N. J., Plenge, R. M., Weinblatt, M. E., Shadick, N. A. and Reich, D. (2006). Principal components analysis corrects for stratification in genomewide association studies. Nature Genetics 38 904–909.

Price, A. L., Zaitlen, N. A., Reich, D. and Patterson, N. (2010). New approaches to population stratification in genome-wide association studies. Nature Reviews Genetics 11 459–463.

Pritchard, J. K. and Przeworski, M. (2001). Linkage disequilibrium in humans: models and data. The American Journal of Human Genetics 69 1–14.

Sabatti, C. (2013). Multivariate linear models for GWAS. In Advances in Statistical Bioinformatics (K.-A. Do, Z. S. Qin and M. Vannucci, eds.) 188–207. Cambridge University Press.

Seaman, S. and Müller-Myhsok, B. (2005). Rapid simulation of P values for product methods and multiple-testing adjustment in association studies. The American Journal of Human Genetics 76 399–408.

Servin, B. and Stephens, M. (2007). Imputation-based analysis of association studies: candidate regions and quantitative traits. PLoS Genetics 3 e114.

Smedley, D., Haider, S., Durinck, S., Pandini, L., Provero, P., Allen, J., Arnaiz, O., Awedh, M. H., Baldock, R., Barbiera, G. et al. (2015). The BioMart community portal: an innovative alternative to large, centralized data repositories. Nucleic Acids Research 43 W589–W598.

Stephens, M. (2013). A unified framework for association analysis with multiple related phenotypes. PLoS ONE 8 e65245.

Stephens, M. (2016). False discovery rates: a new deal. Biostatistics.

Sweeting, T. J. (1986). On a converse to Scheffe’s Theorem. The Annals of Statistics 14 1252–1256.

Vilhjalmsson, B., Yang, J., Finucane, H. K., Gusev, A., Lindstrom, S., Ripke, S., Genovese, G., Loh, P.-R., Bhatia, G., Do, R. et al. (2015). Modeling linkage disequilibrium increases accuracy of polygenic risk scores. The American Journal of Human Genetics 97 576–592.

Wakefield, J. (2009). Bayes factors for genome-wide association studies: comparison with P-values. Genetic Epidemiology 33 79–86.

Wang, K., Li, M. and Hakonarson, H. (2010). Analysing biological pathways in genomewide association studies. Nature Reviews Genetics 11 843–854.

Wen, X. and Stephens, M. (2010). Using linear predictors to impute allele frequencies from summary or pooled genotype data. The Annals of Applied Statistics 4 1158–1182.

Wen, X. and Stephens, M. (2014). Bayesian methods for genetic association analysis with heterogeneous subgroups: From meta-analyses to gene–environment interactions. The Annals of Applied Statistics 8 176–203.

Wood, A. R., Esko, T., Yang, J., Vedantam, S., Pers, T. H., Gustafsson, S., Chu, A. Y., Estrada, K., Luan, J., Kutalik, Z. et al. (2014). Defining the role of common variation in the genomic and biological architecture of adult human height. Nature Genetics 46 1173–1186.

Yang, J., Benyamin, B., McEvoy, B. P., Gordon, S., Henders, A. K., Nyholt, D. R., Madden, P. A., Heath, A. C., Martin, N. G., Montgomery, G. W. et al. (2010). Common SNPs explain a large proportion of the heritability for human height. Nature Genetics 42 565–569.

Yang, J., Manolio, T. A., Pasquale, L. R., Boerwinkle, E., Caporaso, N., Cunningham, J. M., de Andrade, M., Feenstra, B., Feingold, E., Hayes, M. G. et al. (2011). Genome partitioning of genetic variation for complex traits using common SNPs. Nature Genetics 43 519–525.

Yang, J., Ferreira, T., Morris, A. P., Medland, S. E., Madden, P. A., Heath, A. C., Martin, N. G., Montgomery, G. W., Weedon, M. N., Loos, R. J. et al. (2012). Conditional and joint multiple-SNP analysis of GWAS summary statistics identifies additional variants influencing complex traits. Nature Genetics 44 369–375.

Zhang, H., Wheeler, W., Hyland, P. L., Yang, Y., Shi, J., Chatterjee, N. and Yu, K. (2016). A powerful procedure for pathway-based meta-analysis using summary statistics identifies 43 pathways associated with type II diabetes in European populations. PLoS Genetics 12 1–28.

Zhou, X., Carbonetto, P. and Stephens, M. (2013). Polygenic modeling with Bayesian sparse linear mixed models. PLoS Genetics 9 e1003264.

Zhou, X. and Stephens, M. (2012). Genome-wide efficient mixed-model analysis for association studies. Nature Genetics 44 821–824.

